# Neural dynamics of parametrically modulated human mechanical pain determined using whole-brain quantitative perfusion imaging

**DOI:** 10.1101/2021.03.25.436356

**Authors:** A.R. Segerdahl, Y. Kong, I. Ho, I Tracey

## Abstract

Arterial spin labelling (ASL) FMRI is a powerful tool to noninvasively image tonic and ongoing pain states in both healthy participants and patients. We used ASL to image the neural correlates of extended, parametrically modulated mechanical pain in healthy human participants. The aims of this study were to: i) assess if force-calibrated pin-prick probes could safely and robustly evoke tonic mechanical pain; ii) determine the neural correlates of the parametric changes in both the “force” of the stimulus and the “intensity” of the perception that this elicits using ASL; and iii) provide an initial assessment of the capacity for ALFF to differentiate painful versus non-painful tonic stimuli based on changes in the dynamics of the evoked signal. Our data confirm that it is possible to employ a stimulus force-locked design to induce robust, well maintained ongoing mechanical pain and to observe significant changes in rCBF relative to underlying component processes such as monitoring graded changes in the force applied to the skin (dACC, aMCC, pMCC, PCC, SI, SII, putamen, thalamus and the insula (anterior and posterior subsections); ipsilateral amygdala and hypothalamus; and the contralateral DLPFC) and tracking changes in the perceived intensity of the experience (: bilateral dACC, aMCC, pMCC, PCC, thalamus, SII and the cerebellum; and contralateral SI, insula (including the dpIns). Further exploration of the data using analyses targeting the spectral frequency aspects of the rCBF signal observed reveals that a collection of regions (e.g. the contralateral VLPFC, inferior frontal gyrus, insula (anterior, mid and posterior subsections), SII, putamen, OFC, amygdala, and the hippocampus) exhibit unique perfusion dynamics during extended painful stimulation compared to non-painful ‘touch’. Results from this study provide further validation for the application of ASL to image experimental pain in healthy human subjects while interrogation of the data offers unique insight into the dynamic signal changes underlying the perception of a tonic mechanical pain experience.

## Introduction

Imaging tonic, slowly fluctuating and spontaneous pain states in healthy controls or in patients is an emerging focus within the pain neuroimaging community. This is because of the importance of ongoing pain as a key symptom in persistent chronic pain states. Understanding the brain networks subserving this important feature of pain has not been easy. In part, this is because of the difficulty in inducing a continuously active pain state in humans that is experimentally well controlled, does not habituate, is non-invasive and poses minimal risk of skin damage. Another key obstacle is that painful experiences that extend for longer than a few minutes (i.e. non-acute pain) are notoriously difficult to image in a robust and reliable way using standard fMRI tools, such as Blood Oxygen Level Dependent (BOLD) imaging. Specifically, spontaneous and ongoing pain is by definition a non-steady state. It can therefore evade accurate detection with BOLD sequences because the low-frequency drift inherent to this imaging method precludes accurate measurement of neural activity over long durations (Wang et al., 2003). Recent advances in arterial spin labelling (ASL) quantitative perfusion imaging methods minimize some of these concerns and as a result these tools have become more commonplace in both research and clinical FMRI settings (Alsop et al., 2014; (Chappell M et al. 2018). In the pain neuroimaging community, a growing body of work has further optimized the capacity of ASL to image tonic and ongoing pain experiences in a statistically robust and reliable way (Loggia M, Segerdahl AR et al. 2019).

The aim of the current study is to implement a single-post-labelling-delay (PLD) pCASL sequence to image an extended acute mechanical pain state in healthy human subjects where the nociceptive origin of the pain state is well characterized. Previous work interrogating the peripheral origin of continuous mechanical pain shows that it involves both C- and A- peripheral nerves and activity depends on the force, duration and surface area stimulated (Kenshalo et al. 1979; Adriaensen H et al. 1984; Cooper et al. 1993; Greenspan and McGills, 1991; Cervero F. et al. 1988; (Andrew D. and Greenspan J.D. 1999; Slugg et al. 2000).

Forward translation work by Andrew and Greenspan showed that the nociceptor discharge profiles recorded in rodents align with the subjective pain experiences reported by healthy control participants undergoing an identical experimental procedure (Andrew & Greenspan, 1999). The primary finding here was that the perceptual experience of sustained mechanical pain was linked to the stimulus-response properties of the slow-adapting A-fibres; namely that they are activated by nociceptive input to the skin, do not readily adapt with continuous stimulation over the 2-minute block, and show a monotonic increase in firing-rate as the stimulus intensity increases (Andrew & Greenspan 1999).

We aimed to extend these findings further by adapting them to an FMRI setting to identify the cortical correlates linked to the graded changes in the magnitude of force applied to the subject’s hand; and to the changes in perception of pain perceived during each extended-acute mechanical stimulation.

We hypothesized that it will be possible to induce graded changes in mechanical pain for each 5-minute stimulus block using force-calibrated pin-prick probes adapted to an ASL FMRI setting. Additionally, we hypothesized that cortical regions well known from BOLD FMRI investigations of acute pain will also be activated during this type of extended pain experience; with regions such as the thalamus, divisions of the insula and SII showing a significant correlation between rCBF activation and both the magnitude of force applied and the intensity of the pain reported.

Further, we aimed to explore the dynamics of the pain-induced rCBF time courses by assessing changes in low frequency oscillations within active voxels. To do this, we compared the power spectrum within a low-frequency band (<0.1Hz) for each voxel in the brain relative to the global mean power spectrum across the whole brain (Zuo et al., 2009; Zang et al., 2007). The basic principle here is that individual neurons have an intrinsic capacity to oscillate at a range of frequencies. It follows that ensembles of neurons that are synchronized in time form a network, which enables the brain to generate perceptual experiences (Buzsaki & Draguhn, 2004). Visualising the spatial distributions of oscillatory activity can provide insight into neural dynamics underlying a given brain state – either at rest or during a task. Previous work suggests that it is possible to take advantage of methods like this to improve the interpretability of both resting state and task-evoked activity changes that are not captured by standard GLM analysis approaches. An extensive literature exists about the utility of measuring low frequency oscillations across a range of experimental scenarios in both healthy participants (Cauda et al. 2010; Baliki et al. 2011; Baria et al. 2011; Rogachov et al. 2016) and in different patients groups (Baliki et al. 2011; 2014; Hodkinson et al. 2016; Alshelh et al. 2016; 2018) – with more recent work highlighting how these methods may be sensitive to gender-related differences in cortical processing of pain (Hong et al. 2013; Rogachov et al. 2018). We investigated low-frequency osciallations using a common ALFF tool in the current study to investigate how this might also be observable during an extended acute mechanical pain experience and if this might be a way to further interrogate the meaningfulness of the pain related rCBF changes observed.

## Methods

Experiments were performed on 18 healthy subjects (10 female; right handed; age: 18-40yrs). Subjects were asked to partake in the psychophysics protocol explained below whilst being scanned using a Siemens 3T Verio scanner. No one was on medication likely to interfere with the pain responsiveness. All subjects gave informed consent and the Oxfordshire Clinical Ethics Committee approved this study.

### Stimuli

Mechanical probes (MRC, Germany) calibrated to give a constant force (64mN, 256mN and 512mN) were each applied to the skin separating the thumb and first finger on the subject’s right hand for a period of 5 minutes. A homemade plastic handle was used to comfortably expose the stimulus site over the duration of the experiment (Figure 1a).

**Figure 1.**
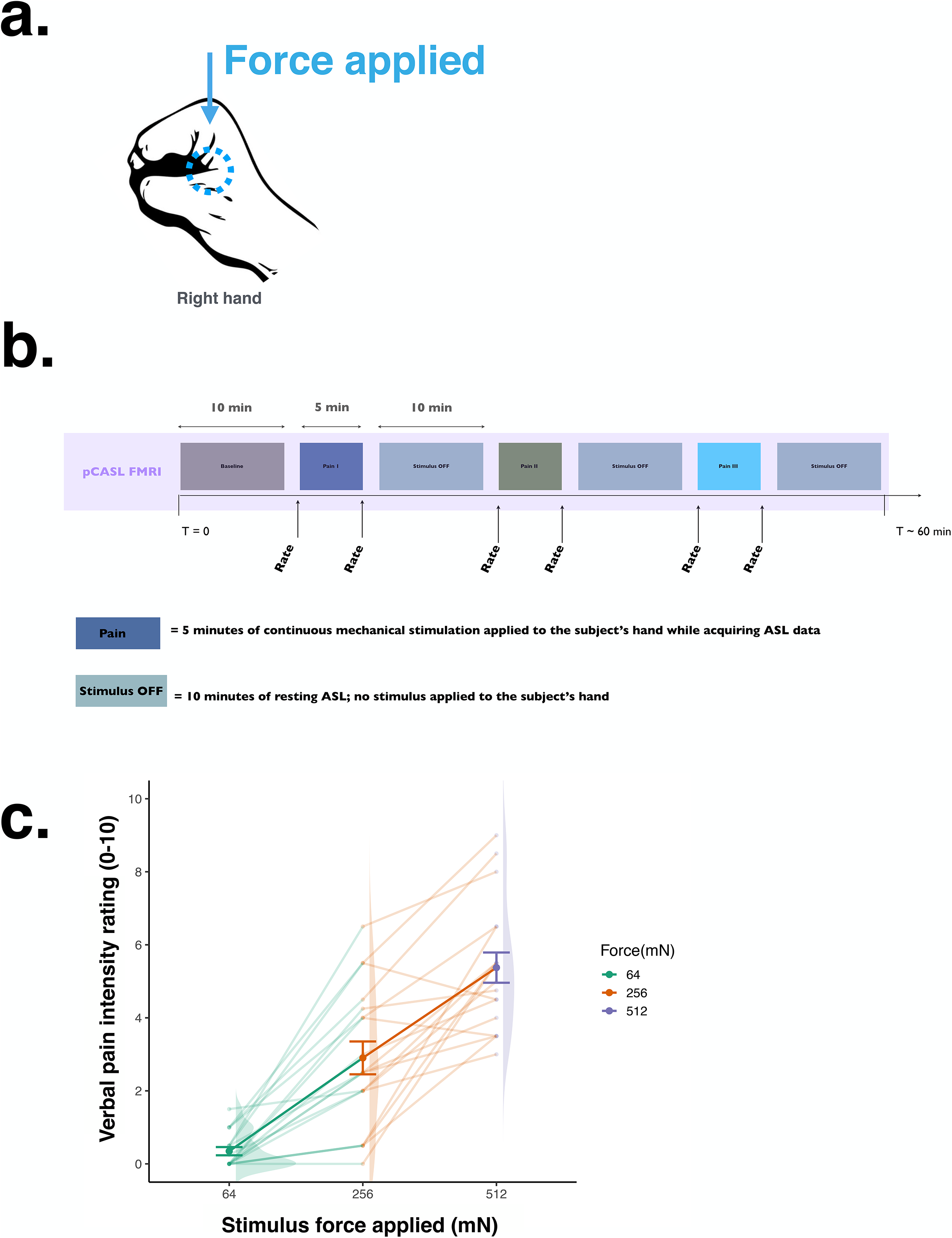
Paradigm design. a) A schematic showing the stimulation site on the subject’s right hand. b) A timeline of the experimental paradigm and scan protocol used in the study. The 10-minute rest blocks are displayed in grey. Each 5-minute stimulus block is coloured. Each force applied is represented by a different colour. The stimulus order was pseudo-randomised across subjects. Verbal pain intensity and unpleasantness ratings were obtained 30-seconds after the stimulus onset using the Numerical Rating Scale (NRS: 0 = no pain; 10 = worst pain imaginable). This immediately preceded the acquisition of the first imaging volume. A second pain rating was obtained immediately after the last imaging volume was acquired for each stimulus condition. c) Group mean pain intensity (colour) ratings for each stimulus condition (green = 64mN force, Lp; orange = 256mN force,Mp; purple = 512mN force, Hp). Data are representative of all 18 subjects that completed the scan session. Significant differences in reported pain perception across each stimulus condition are marked with a * (n=18; one-way ANOVA, p<0.001). Error bars represent the standard error of the mean (± s.e.m.).

### Psychophysics

Subjects were told they would receive a series of stimulations that would last for a few minutes but were not told the exact duration of the stimulus block. Mechanical stimuli were applied to the subject’s right hand by placing the probe into a stabilizing apparatus that securely held it at a constant force on the stimulus site for the duration of the 5-minute stimulus block. The stimulus order was pseudo-randomized across subjects to control for potential confound effects of stimulus order, sensitization and habituation. A 10-minute rest block (no stimulus present) was recorded after each stimulus block. Here, subjects were asked to maintain the position of their hand but in the absence of a stimulus probe. See Figure 1b for an illustration of the paradigm used. During the stimulus, subjects were asked to keep their eyes closed and to focus on the stimulation being received. Subjects rated the stimulus intensity verbally using an 11-point numerical rating scale (NRS; 0= no pain; 10= worst pain imaginable) every 30 seconds. During the imaging session, verbal ratings were taken 30-seconds after the stimulus onset but before the 5-minute ASL scan was acquired. A second verbal rating was taken immediately after the last imaging volume was acquired. Verbal pain scores were then averaged for each subject to generate an overall pain rating for each stimulus block scanned.

### FMRI Pain Protocol

Once the subject was placed in the scanner, a ten-minute resting baseline scan was acquired. Immediately after, mechanical stimuli were delivered as discussed previously. Ratings were reported verbally after the first 30-seconds of stimuli and then after the 5-minute scan. The 30-second ‘rating phases’ that flanked the five minute ‘pain scan’ were not scanned as discussed above. Ten-minute rest blocks following each pain block were also scanned. Pain intensity and unpleasantness scores were taken at the end of each rest block to monitor any residual pain effects. Subjects were asked to remain still with their eyes oriented to a standard fixation cross throughout the entire scan session.

### Single-PLD pCASL FMRI Parameters

All subjects were scanned with a pseudo-continuous ASL sequence (Dai et al. 2008) using gradient-echo EPI readout (TR=3.75s, TE=13ms, 6/8 k-space). Twenty-six axial slices in ascending order (3×3×4.5mm voxels, 0.5mm inter-slice gap) were used for each subject to provide coverage of the whole brain. The labelling offset was defined 10cm inferior to the center of the 26 slices. A 90° pre-saturation pulse was applied before the labelling pulses. Labelling duration was 1.4s and a single post-label delay time of 1.0s was used. All subjects were scanned using a Siemens 3T Verio system fitted with a 32-channel head coil.

### ASL Analysis

#### I. Calculation of whole brain relative perfusion (rCBF) maps

ASL data were analyzed using the FMRIB Software Library (FSL) (Smith et al. 2004). At the first level, the following pre-processing steps were carried out: brain extraction with Brain Extraction Tool (BET) (Smith et al., 2002), motion correction with MCFLIRT (Jenkinson et al., 2002) and spatial smoothing with a Gaussian kernel of full-width-half-maximum = 5mm. Intensity normalization was carried out with a single scaling factor and high-pass temporal filtering performed with a Gaussian-weighted least squares straight line fit and high-pass cut off filter of 320s. Data were de-noised of non-physiological artifacts through visual inspection of independent component maps generated using MELODIC (ICA citation; Kelly, et al. 2010). Registration of subject’s functional datasets was completed in three stages: 1) a single-slice EPI image from one subject was used as an initial structural image to register all subject’s main high-resolution structural images into a standard orientation. Subsequently, subject’s functional datasets were co-registered onto their high-resolution structural images and finally to a standard MNI152 T1-weighted brain using a nonlinear transformation (FNIRT, FSL) (Smith et al. 2004).

Statistical analysis was performed with the use of FMRIB’s Expert Analysis Tool (FEAT) (Smith et al., 2002). At the first level, each subject’s perfusion timeseries was generated by fitting a model of the relevant ‘tag’ and ‘control’ images to the denoised 4D dataset. Each voxel was fit to the perfusion model to yield a resultant parameter estimate (PE) image that was proportional to localized blood flow. To determine the change in rCBF associated with the pain stimuli, a model of each stimulus force (e.g. 64mN, 256mN and 512mN) was fit to each subject’s perfusion time series using a ‘fixed effects’ second level analysis. At the top level, we implemented FSL’s tool for nonparametric permutation inference testing of neuroimaging data called Randomise to identify rCBF changes (TFCE: threshold-free cluster enhancement; familywise error (FWE) corrected p-values <0.05) (Winkler AM et al. 2014).

To describe regions that track graded changes in the amount of nociceptive input applied, the forces used to elicit the “low”, “medium” and “high” pain states (e.g. 64mN, 256mN, and 512mN) were modeled as regressors of interest in a repeated measures ANOVA design (Randomise, TFCE; FWE-corrected p<0.05). The results from this analysis reveal those regions within which the extent of hyper-perfusion is correlated to the magnitude of force applied to the subject’s hands (Figure 2a).

**Figure 2:**
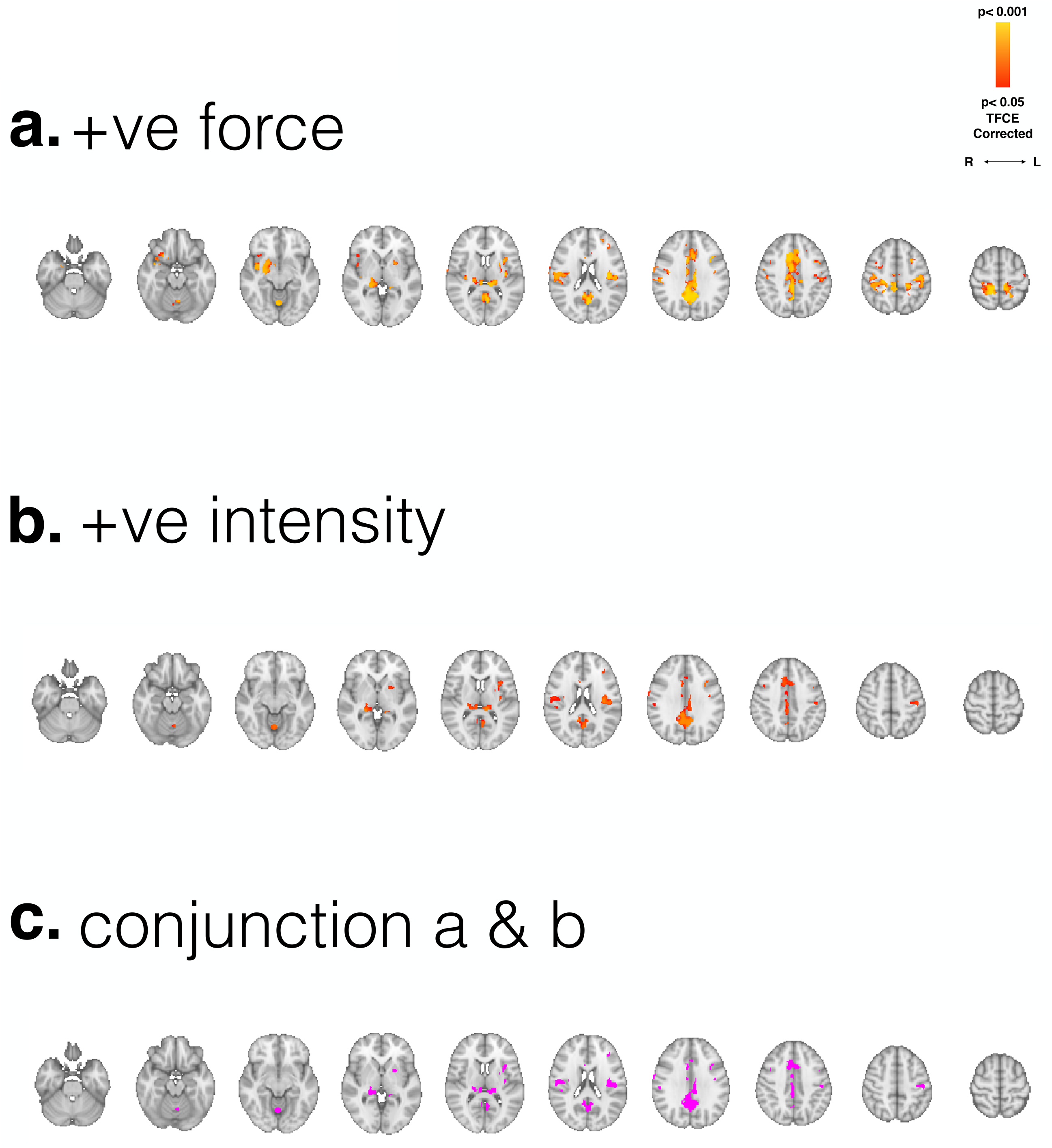
Whole-brain rCBF changes during pain. a) Regional rCBF changes during mechanical stimulation that shows a significant correlation with a graded increase in stimulus force (i.e. 64mN < 256mN < 512mN) are displayed. Voxels with supra-threshold activation are shown in red and are overlaid on the MNI152 standard template brain (n=18, Randomise TFCE FWE-corrected p<0.05). b) Voxels within which a significant positive correlation between rCBF and the reported pain intensity ratings elicited by increasing levels of mechanical stimulation are displayed in red and are overlaid on the MNI152 standard template brain (n=18, Randomise TFCE FWE-corrected p<0.05). c) A conjunction analysis was used to identify voxels within which co-activation was observed for both force (a) and intensity (b) tracking across each force applied (purple). Axial slices correspond to the plane indicated in the sagittal slice (MNI z coordinates: −32 to 70). Radiological convention is used (L: Left; R: Right).

To further characterize the function of these pain-related regions, we completed a second regression analysis to interrogate within which regions the extent of hyper-perfusion observed during mechanical stimulation is correlated to the magnitude of pain intensity experienced by the subjects (Figure 2b). Here, the verbal pain intensity ratings reported for each stimulus condition were included as a regressor of interest in a repeated measures ANOVA design (Randomise, TFCE; FWE-corrected p<0.05).

To explore how the ALFF index changes across the whole brain during ongoing pain, we investigated the presence of altered ALFF index when comparing “pain” versus “non-painful touch” in FEAT (Randomise, TFCE; FWE-corrected p<0.05). Briefly, the ALFF index is calculated by averaging the square root of the power spectrum within a low-frequency band (<0.1Hz) for each voxel. This value is then normalized relative to the global mean ALFF value across the brain (Zuo et al., 2009; Zang et al., 2007). This analysis shows voxels within which there is a significant alteration in the amplitude of rCBF fluctuations occurring during the painful stimulation compared to during non-painful stimulation (Figure 3a). To further investigate the meaningfulness of these changes, a conjunction analysis was used to determine if regions that show a significant alteration in rCBF dynamics, as determined with ALFF, also are observed to have a significant correlation between rCBF and the magnitude of the force applied (Figure 3b). Results from this exploration may illuminate regions within which the functional dynamics (as measured by ALFF) might be linked to the nociceptive force-tracking capacities of a core set of regions during the experience of extended mechanical pain.

**Figure 3:**
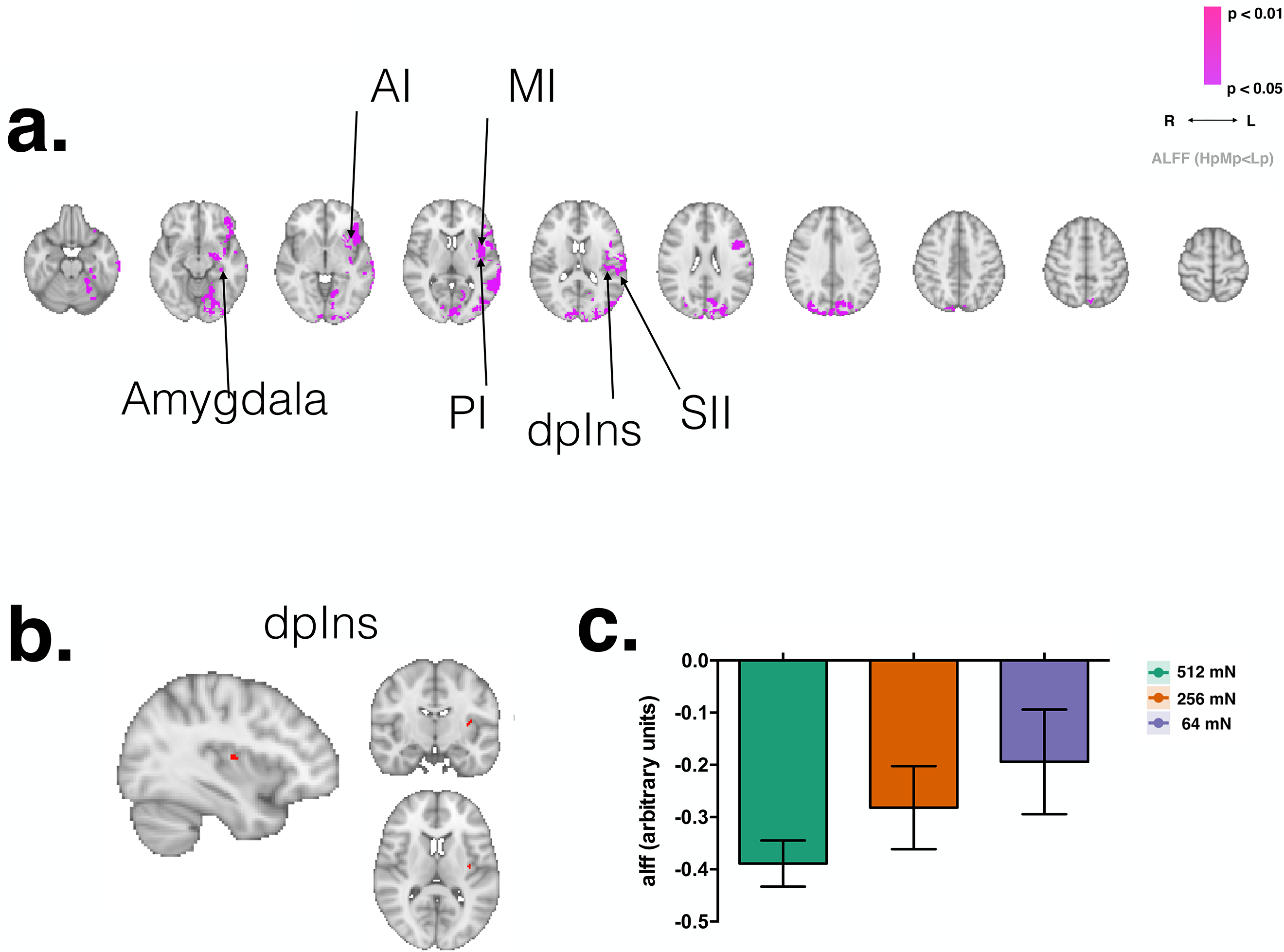
Whole-brain amplitude of low-frequency fluctuations (ALFF) changes during painful versus non-painful touch. *a)* Voxels showing a significant decrease in ALFF index during painful versus non-painful stimulation are displayed in purple and are overlaid on the MNI152 standard template brain (n=18, Randomise TFCE FWE-corrected p<0.05). b) A conjunction analysis that was used to identify voxels within which a change in ALFF index overlaps with rCBF changes that correlate with the force of the mechanical pin-prick probes applied to the hand is displayed in Figure 3b (red). Axial slices correspond to the plane indicated in the sagittal slice (MNI z coordinates: −32 to 70). Radiological convention is used (L: Left; R: Right).

## Results

### 1. Psychophysics

Figure 1c shows the mean pain intensity ratings for the three mechanical forces used in this study (64mN, 256mN, and 512mN). A repeated measures ANOVA showed that the group mean pain intensity ratings differed significantly between the forces applied to the participants [F(1.98, 33.12)=68.90, p<0.001]. Post hoc tests using Tukey’s correction for multiple comparisons revealed that the pain intensity ratings increased by an average of 2.556 when the force was increased from 64mN to 256mN (95% confidence interval [Cl] = 1.492, 3.619; p<0.0001). The pain ratings increased again by an average of 2.472 when the force was increased from 256mN to 512mN (95% confidence interval [Cl] = 1.288, 3.657; p<0.001).

### 2. ASL FMRI

#### i. Cortical regions tracking graded changes in the stimulus magnitude

Cortical regions within which the extent of hyper-perfusion is correlated to the magnitude of force applied to the subject’s hands are displayed in Figure 2a. Regions in colour represent voxels with supra-threshold z-statistics generated from the group level FEAT analysis described previously. Significant hyper-perfusion related to a graded increase in stimulus force (from 64mN to 512mN) applied to the subject’s hands occurred in regions such as: bilateral supplementary motor area (SMA), cingulate cortex (including the dorsal anterior cingulate: dACC; anterior mid-cingulate: aMCC; posterior mid-cingulate: pMCC; and posterior cingulate: PCC subsections), primary somatosensory cortex (SI), secondary somatosensory cortex (SII), putamen, thalamus and insula (anterior and posterior subsections including the dorsal posterior insula (dpIns)), ipsilateral (right) amygdala, hypothalamus and the contralateral dorsal lateral prefrontal cortex (DLPFC).

#### ii. Neural correlates of ongoing mechanical stimulation that track changes in perceived pain intensity

Regions recruited during the experience of tonic pain that show a significant change in rCBF related to the increase in pain intensity perceived by subjects are shown in Figure 2b. Significant hyper-perfusion related to the reported pain intensity scores occurred in regions such as the: bilateral dACC, aMCC, pMCC, PCC, thalamus, SII and the cerebellum; and contralateral SI, insula (anterior and dpIns), and the putamen. For display purposes, a conjunction analysis was used to identify voxels within which co-activation was observed for both force (a) and intensity (b) tracking across each force applied (Figure 2c). Regions identified include the: bilateral bilateral dACC, aMCC, pMCC, PCC, thalamus, SII and the cerebellum; and contralateral SI and the dpIns. The direct comparison between “intensity” versus “force” did not reveal regional mismatches in activation that survived correction (Randomise, TFCE; FWE-corrected p<0.05). For reference, the uncorrected statistical maps are provided for inspection (Supplementary Figure 1). Briefly, “Rate>Force” yields CBF changes in hippocampus, cerebellum, secondary somatosensory cortex (SII), dorsal lateral prefrontal cortex (DLPFC) and inferior frontal gyrus; “Force > Rate” produces CBF changes in the primary somatosensory cortex (SI), precuneus, cingulate (posterior, mid and anterior subsections), dorsal medial prefrontal cortex (DMPFC), ventral medial prefrontal cortex (VMPFC), insula (posterior and anterior subsections) and the amygdala.

#### iii. Investigation of the dynamic changes in rCBF during pain using frequency domain analysis

Figure 3 shows the results of the ALFF analysis. Regions in purple represent voxels with significant decreases in amplitude of CBF signal fluctuation during pain compared to non-painful stimulation (HPMP vs. NP). These regions include: contralateral ventral lateral prefrontal cortex (VLPFC), inferior frontal gyrus, insula (anterior, mid and posterior subsections), SII, putamen, OFC, amygdala, and the hippocampus. No significant increases in amplitude of the CBF signal fluctuation during pain were observed. For display purposes, we overlaid the perfusion maps in Figure 2c (conjunction between FORCE x RATE) and in Figure 3a (ALFF) to identify voxels within which graded changes in both stimulus force and perception were co-localized to areas showing a decrease in CBF signal amplitude during tonic stimulation (Figure 3b). The region that showed this co-localization was the dorsal subsection of the posterior insula (dpIns). The group mean ALFF values extracted from this region are displayed in Figure 3c.

## Discussion

The primary aim of this investigation was to image the neural correlates of ongoing, parametrically modulated mechanical pain in healthy human subjects using a single-TI ASL imaging protocol and standard FSL tools. Our data confirm that it is possible to employ a stimulus force-locked design to induce robust and well maintained ongoing mechanical pain. We can observe significant changes in rCBF related to the underlying component neural processes such as monitoring graded changes in the nociceptive input applied to the skin and tracking changes in the perceived intensity of the experience. Further exploration of the data using analyses targeting the spectral frequency aspects of the rCBF signal observed reveals that a broad set of regions (e.g. the contralateral VLPFC, inferior frontal gyrus, insula (anterior, mid and posterior subsections), SII, putamen, OFC, amygdala, and the hippocampus) exhibit altered perfusion dynamics during extended painful stimulation compared to non-painful ‘touch’. Results from this study provide further validation for the application of ASL to image experimental pain in healthy human subjects while interrogation of the data offers insight into the neural oscillations underlying the construction of an extended mechanical pain experience.

### Neural correlates of extended acute mechanical pain

A key interest of this work is to better understand the neurophysiology of brain processes underlying pain experiences that last for more than a few minutes. In the current study, we employed a single-TI ASL method to image 5-minute blocks of continuous noxious mechanical stimulation applied to the subject’s hand. The benefit of this approach is that it allows for more accurate comparison with other investigations of extended acute and some types of clinical pain that persist on similar timescales.

It is well-accepted that no one region of the brain encodes pain alone. Instead, the experience of pain arises from dynamic processing within a network of brain regions that collectively encodes its key underlying features (e.g. stimulus intensity, location, emotional affect, etc.) and integrates this with other core attentional, mnemonic and cognitive processes. An extensive literature exists supporting this concept (Melzack and Casey, 1968; Bornhovd et al., 2002; Coghill RC et al., 1999; Derbyshire et al., 1997; 1998; Tracey I. 2005; 2007; 2015; Wager T. et al., 2013; Garcia-Larrea, 2013; Kuyci & Davis, 2015; Atlas et al. 2014; Wiech K. et al. 2014; 2016). However, most of this knowledge arises from studies in acute pain where the experience is very brief and phasic. We need to also interrogate how the experience of tonic pain arises from complex spino-cortical network dynamics, but to do this it is necessary to identify which regions are coding key features of an extended mechanical pain experience (e.g. stimulus magnitude, perceived intensity).

Here, we adapted a force-locked parametric pain stimulus design validated previously in animals and humans (Cevero and Handwerker 1988; Woolf and King 1987; Andrew and Greenspan 1999). This method benefits from being able to reliably and robustly induce extended mechanical pain at varying levels of intensity (e.g. light non-painful touch versus moderate to high pain) with minimal risk of skin damage or the incidence of habituation during stimulation. While it is not possible to dissect the cortical activation maps presented here based on the extent of nociceptor subtype involvement; previous electrophysiological work in animals using similar methods have characterized the nociceptive components underlying the perception of extended mechanical pain. Results from these studies show that multiple nociceptor subtypes are involved. These include: i) an initial phasic C- and rapidly-adapting A-fibre component that is excited at pain threshold and has a relatively slow (< 1 HZ) firing frequency; and ii) a sustained higher frequency (>5Hz) tonic discharge from a distinct subtype of A-fibres that are slowly-adapting (Kenshalo et al. 1979; Cooper et al. 1993; Greenspan and McGills, 1991; Cervero F. et al. 1988; Andrew D. and Greenspan J.D. 1999; Slugg et al. 2000).

Using this method, it was possible to determine pain-related brain activity linked to: i) parametric changes in the magnitude of force applied to the subject’s hand (i.e. nociception); ii) graded changes in the intensity of the pain experienced by the subjects (i.e. subjective perception); and iii) alterations in the rCBF dynamic time courses (ALFF) that may reflect pain related neurophysiological features relative to the innocuous touch condition; these would otherwise not necessarily be observable with a standard GLM analysis as has been shown previously with BOLD FMRI studies (Cauda et al. 2010; Baliki et al. 2011; Baria et al. 2011; Rogachov et al. 2016). We attempted to identify what is independently linked to these outcome measures using standard covariance analyses. The data show that within a number of cortical regions, the extent of rCBF changes scale parametrically with the magnitude of force applied to the subject’s hand. These regions include: bilateral SMA, dACC, aMCC, pMCC, PCC, SI, SII, putamen, thalamus and the insula (anterior and posterior subsections and dpIns); ipsilateral amygdala and hypothalamus; and the contralateral DLPFC. These data align with previous parametric pain studies that demonstrate a relationship between the stimulus magnitude and the extent of metabolic signal change observed in key brain regions that are known to be involved in coding aspects of nociception, arousal and attention. These regions include: the: cerebellum, putamen, thalamus, insula (anterior and posterior subsections), cingulate (anterior and posterior subsections), SI, SII, prefrontal and inferior parietal cortices [acute pain imaged with BOLD FMRI: Buchel et al., 2002; Bornhovd et al., 2002; phasic pain imaged with PET: Coghill et al., 1999; Derbyshire et al., 1997; Casey et al. 1996; Tolle et al., 1999; and tonic pain with ASL Owen DG et al. 2008; 2012; 2010; Frölich M.A. et al. 2012; Lin R. et al. 2017; Howard MA et al. 2011;2012; Thunberg et al. 2005; Stohler CS et al. 1999; Svensson PS et al. 1997)].

The measured activity in the following regions positively correlated with perceived pain intensity: bilateral dACC, aMCC, pMCC, PCC, thalamus, SII and the cerebellum; and contralateral SI, insula (anterior and dpIns), and the putamen. One interpretation of these data is that an ongoing experience of pain (i.e. a composite of both the unabated noxious stimulation applied to the skin plus the persistent perceptual awareness of a constant experience of pain related to that stimulus) results from the dynamic activation of primarily higher cortical regions shown previously to be involved in coding: i) features of the stimulus location, intensity, modality (e.g thalamus, posterior insula, SII); ii) the aversive nature of the stimulus (e.g. DMPFC, anterior insula, ACC); and iii) the modulatory circuitry related to attention, expectation, and reappraisal (e.g. DLPFC and hippocampus). These data reveal that the magnitude of activation in each of these regions scales with the severity of the pain experienced.

These data may also reflect the extent to which the engagement of functionally connected regions enables pain-relevant information to be distributed and processed widely across the brain. For example, it is well known that the subdivisions of the insula are both anatomically and functionally connected to a number of brain regions (including the: thalamus, SI, SII, rACC, dACC, pMCC, VLPFC, and DLPFC) (Taylor et al., 2009; Wiech et al., 2014). It is logical that while essential variables underlying the overall pain experience (like the magnitude of force and the intensity of the experience) are tracked by the thalamus and subdivisions of the insula; the overall emergence and maintenance of the pain experience relies on the recruitment and activation of other connected regions such as the cingulate and the prefrontal cortex. For example, the prefrontal cortex has direct anatomical connections to limbic, motor, and sensory areas via brainstem and subcortical pathways; in addition to being densely interconnected with itself and other cortical areas involved in pain perception (Hadjipavlou et al., 2006; Tekin and Cummings 2002; Wiech et al., 2008). As such, the prefrontal cortex provides a core “cognitive control” mechanism that guides how incoming nociceptive input is processed based on the motivations and expectations that have been learned in relation to the stimulus being experienced (Miller and Cohen, 2001; Wiech et al. 2016; Jepma et al. 2018; Seymour et al. 2019).

We also aimed to explore differences in regional activity when there is a mismatch between perception (pain ratings) and nociceptive input (force applied). The direct comparison of the relationship between CBF and these covariates of interest did not resolve significant differences. In part, this is not surprising given how tightly correlated force and ratings are in this experiment; our approach might simply be too underpowered to detect orthogonal effects. However, a key study by Atlas and colleagues (2014) was able to interrogate this using a BOLD FMRI approach by assessing trial-by-trial differences in subjective reports when varying levels of acute noxious thermal stimuli were applied to healthy participants forearms (Atlas et al. 2014). Using a multi-level mediation analysis, the authors were able to identify regions that significantly mediate pain independent of a purely nociceptive mechanism. Specifically, deactivation of core nodes of the DMN – including the hippocampus, parahippocampus and precuneus - were positively correlated with the intensity of heat pain experienced; an effect that is well aligned with what is known about the fundamental properties of this task-negative network and its relationship to pain (Seminowicz & Davis 2007; Loggia et al. 2013). However, in the current study, inspection of the uncorrected maps points shows enhanced hippocampal, SII, inferior frontal gyrus and prefrontal cortex within the comparison of Rate > Force. One possibility is that this is due to active pain facilitation, possibly driven by heightened emotion (e.g. state anxiety). Previous work shows that experimentally manipulating anxiety can enhance the subjective experience of pain even for identical nociceptive input to the body & this is linked to increased activation of the hippocampus (Ploghaus et al. 2001). Enhanced hippocampal activity (along with the medial PFC and cerebellum) has been shown to underlie negative treatment expectancy on opioid analgesia. Here, pain severity increased – even during infusion of remifentanil analgesia – when participants were induced to think that the analgesic was not effective (negative expectancy) (Bingel et al. 2012). Although we are not able to explore this further with the current dataset, it is interesting to note the potential involvement of this region in the perception of tonic pain and how dysfunction here may engage a distinct supraspinal mechanism of hyperalgesia during persistent pain states.

### Pain related changes in CBF dynamics

Presently, this study is not well suited to interrogate rigorously the dynamic interaction between the regions we show to be linked to aspects of the tonic pain state. However, as a first attempt, we employed an ALFF index to observe how pain triggers specific changes to regional activation over time (as measured by changes in oscillatory dynamics) during the extended pain stimulus. The functional relevance of low frequency oscillatory activity related to brain function in ASL data is not well known even though such analyses are commonplace in other electrophysiological investigations like EEG (Laufs et al. 2008; Mantini et al., 2007). In general, it is thought that changes in oscillations of neural activity as observed with FMRI are likely to reflect either a change in ongoing neural activity, changes in cortical excitability or may also reflect non-neural sources including changes in physiology, imaging related noise and artefacts – features which we have attempted to account for within the data preprocessing pipeline (Wise et al., 2004; Cordes et al., 2001; Mitra et al., 1997; Cole et al. 2010; Van Dijk et al. 2012).

Importantly, a few studies have successfully linked alterations in oscillatory dynamics within cortical regions to changes in behavioural states (Duff et al. 2008; Baria et al 2011) and pain (Cauda et al., 2010; Malinen et al., 2010; Baliki et al., 2011). These more recent investigations in pain patients successfully employed frequency analysis to link alterations in oscillatory dynamics of the BOLD FMRI signal to changes in the functional connectivity of key brain regions (Baliki et al., 2011; Cauda et al., 2010; Malinen et al., 2010). At present, no one has investigated how oscillatory dynamics in ASL FMRI time courses reflect meaningful changes in brain function during a parametrically modulated extended pain state in healthy subjects. Instead, it is common to refer to the average signal change (i.e. rCBF, BOLD) during pain versus rest to describe meaningful differences in neural activity. However, it is likely that the dynamics of the signal time course across the brain during sustained activation also reveal meaningful neurophysiological information linked to pain (Peng W et al. 2014; Kucyi A. & Davis K.D. 2016). A recent paper by Lee et al. (2021) supports this by identified a neuroimaging biomarker for sustained experimental and clincal pain by focusing on dynamic changes in brain function (Lee et al. 2021). Here, by using multiple BOLD FMRI datasets these authors employ a rigorous, data-driven approach to build a predictive algorithm for both experimental tonic and clinical pain that is based on whole-brain dynamic connectivity changes. Importantly, these authors show that by comparison, relying on static changes (e.g. static correlation averaged over scan duration) alone had lower predictive accuracy & had poor cross-validation performance (Lee et al. 2021).

In the current study, the data suggests that during the experience of extended mechanical pain (compared to non-painful touch stimulation), a core set of recruited regions not only have increased CBF but their perfusion dynamics appear to become tuned (i.e. CBF signal decreases in amplitude) such that the region is continuously “on” and hyper-perfused in a steady active state. Given the nature of the stimulus paradigm (i.e. tonic mechanical stimulation that subjects do not habituate to), we posit that the nociceptive afferent input arising from the periphery is continuous and this, in turn, produces a sustained cortical response. We explored this by visualizing those regions that were commonly activated in the assessment of parametric changes in stimulus force (Figure 2a) and pain-induced changes in ALFF (Figure 3a). We observed that the contralateral posterior insula is visible (Figure 3b) - a region shown previously to code physical pain related information about stimulus location and intensity, as well as encoding tonic pain states (Craig, 2014; Evrard et al., 2014; Brooks et al., 2005; Henderson et al., 2007; Baumgartner et al., 2010; Mazzola et al., 2009; Segerdahl et al., 2015; Geuter et al., 2017). One interpretation of this result is that during painful constant stimulation, this region remains in a steady-state of activation throughout the 5-minute stimulation block. Recent work on steady-state functional tasks suggests that continuous activation results in changes in amplitude of the signal within active regions and this can negatively impact the interpretability of functional connectivity assessments if not accounted for (Duff et al. 2018). While the current study is not designed to interrogate this complexity further, future investigations will need to take this into account in order to explore how key nodes, such as sub-sections of the insula, are communicating with other brain regions during ongoing pain; how these functional relationships change over time and across stimulus type; and the extent to which these relationships may become dysfunctional in the context of different chronic pain states.

---

**Supplementary figure 1:**
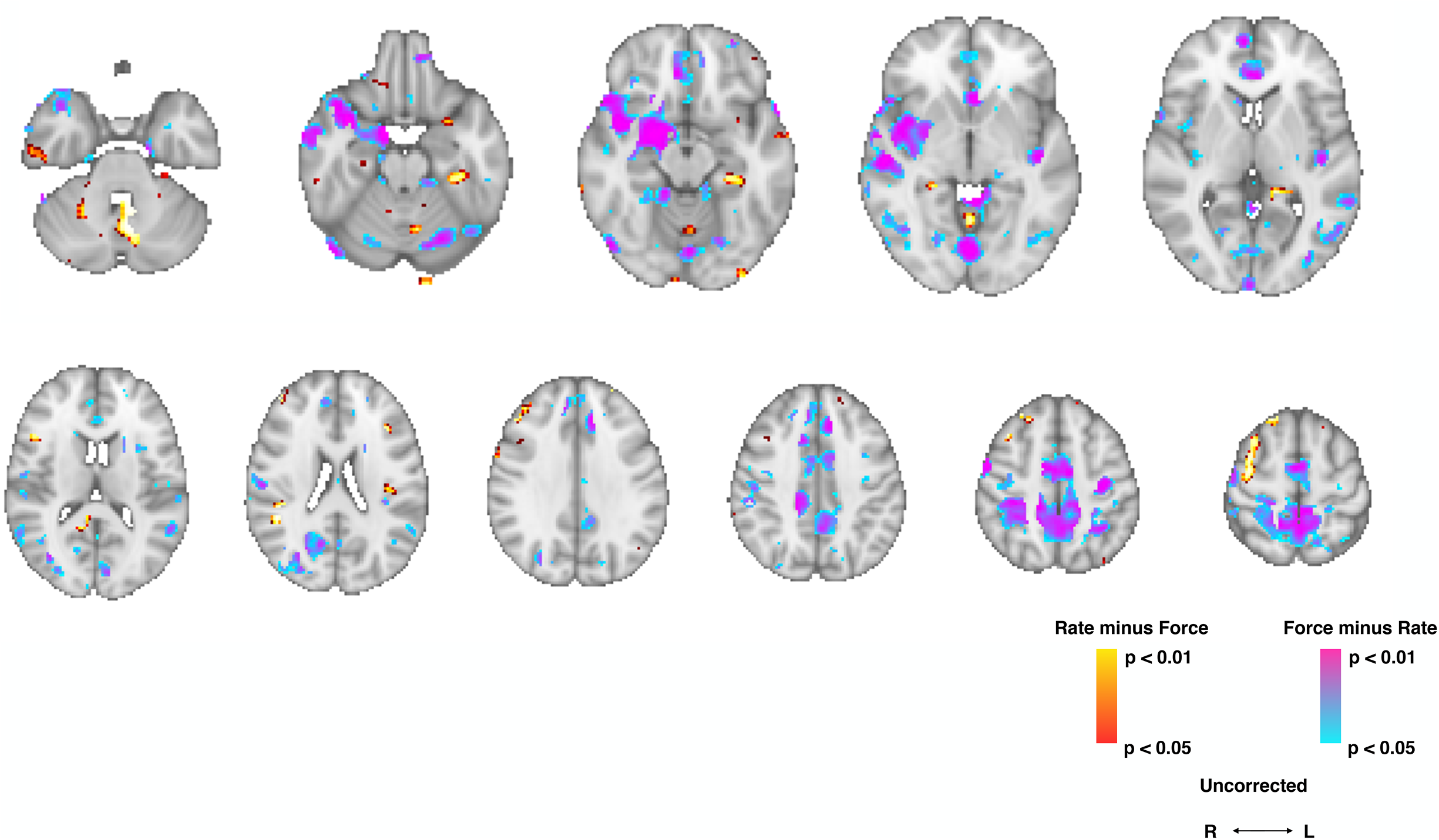
Direct comparison between “intensity” versus “force”. No regional mismatches in activation survived correction (Randomise, TFCE; FWE-corrected p<0.05). For reference, the uncorrected statistical maps are displayed. Briefly, “Rate>Force” is displayed in red (TFCE uncorrected); “Force > Rate” is displayed in purple (TFCE uncorrected). Axial slices correspond to the plane indicated in the sagittal slice (MNI z coordinates: −32 to 70). Radiological convention is used (L: Left; R: Right).

